# The protein interactome of *Escherichia coli* carbohydrate metabolism

**DOI:** 10.1101/2024.03.12.584555

**Authors:** Shomeek Chowdhury, Stephen S. Fong, Peter Uetz

## Abstract

Carbohydrate metabolism is strictly regulated by multiple mechanisms to meet cellular needs (i.e. energy production). Several mechanisms modulate the amount and activity of metabolic enzymes. Here, we investigate how carbohydrate metabolism (CHM) in *E. coli* is regulated by their interaction properties with other proteins and their quantities. We computationally analyze 378 protein-enzyme interactions (PEIs) potentially involved in carbohydrate metabolism. We identified 20 enzymes and 19 interactors that occur at stoichiometries that are highly likely to affect CHM and 174 interactions that are possibly conserved across thousands of bacteria. These PPIs are predicted to be of global importance, including pathogens.

**Author summary:** In systems biology, “big” data are used to reconstruct biological networks which are then investigated in detail to uncover their functional dynamics. Understanding functional dynamics finally helps in identifying biomarkers in the big networks. We apply this concept to the biological network which is a crosstalk of protein-protein interaction network and carbohydrate metabolic pathway in *Escherichia coli* K-12. We investigate if proteins regulate enzymes when they bind them and we predict that protein-enzyme interactions (PEIs) regulate carbohydrate metabolism in *E. coli*.

## Introduction

Carbohydrate metabolism (CHM) is responsible for converting sugars and other compounds into other metabolites and ATP and occurs in all organisms from bacteria to humans [1]. Carbohydrate metabolism includes about a dozen sub-pathways in bacteria and thus can involve more than a hundred metabolic enzymes [1]. In *E. coli*, the focus of this study, CHM involves 13 sub-pathways (**Table 1**, **Fig 3**).

**Table 1.**
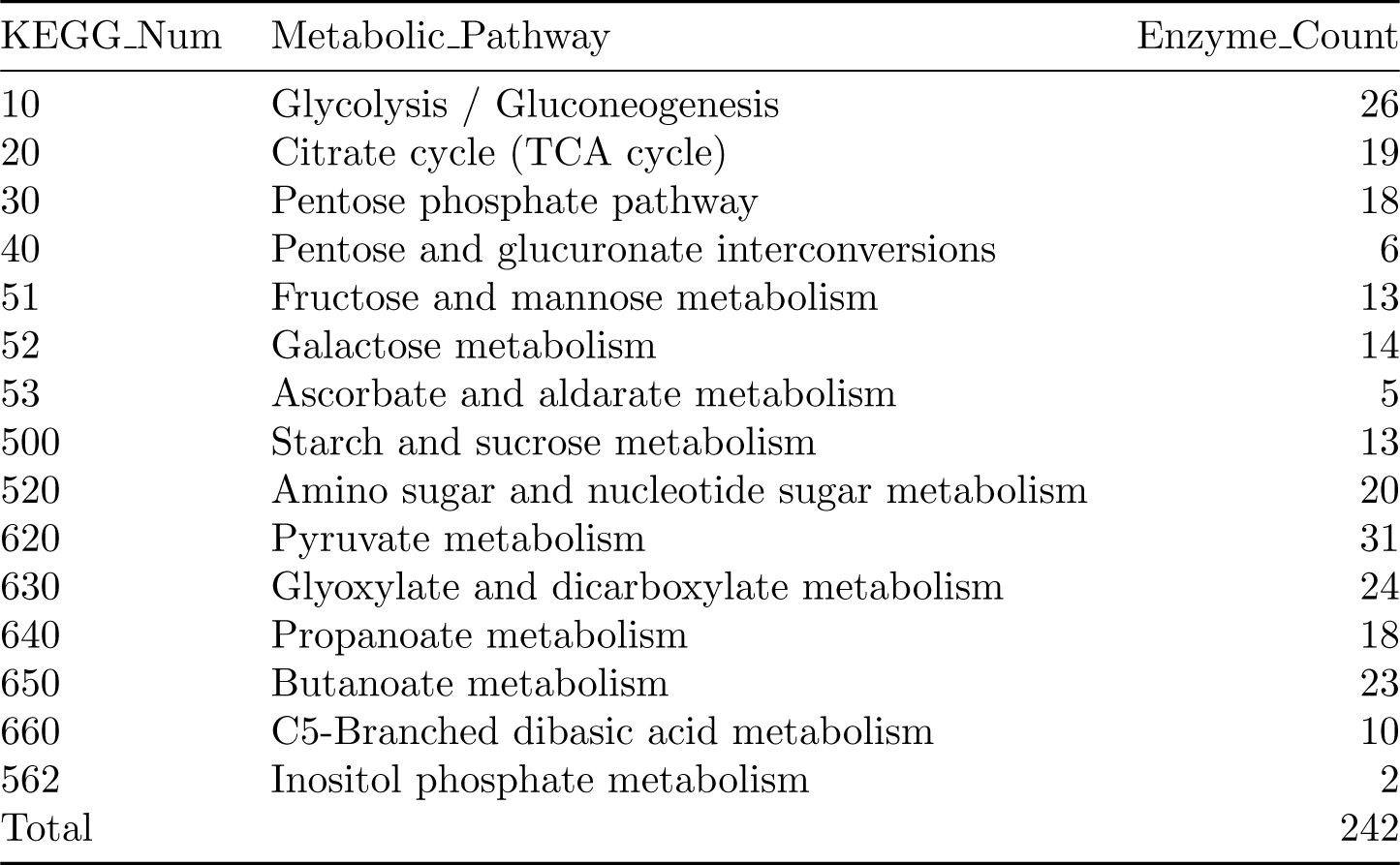
Number of enzymes of *E. coli* carbohydrate metabolism pathways based on KEGG. [2].

Central carbon metabolism is the shortest route for producing energy and biomass in Escherichia coli [3]. In addition, CHM provides the substrates for many other metabolic pathways, such as lipid or amino acid metabolism [2,4].

Given that carbohydrate and energy metabolism in general is critical for cellular survival, it is usually tightly regulated [3]. Different regulatory mechanisms have been identified, including gene expression [5], post-translational modification (PTMs) [6], enzyme localization [7], or allosteric control [8].

Protein-protein interactions (PPIs) are a common mechanism to regulate protein activity. Although numerous enzymes have been shown to be regulated by PPIs, it is difficult to study the impact of PPIs on metabolism on a large scale. For instance, we have demonstrated how several metabolic enzymes in *Escherichia coli* are regulated by PPIs, including enzymes in energy metabolism, carbohydrate metabolism or amino acid metabolism [8–11]. Notably, a single protein such as *Hpr* can both activate and repress enzymes in the same pathway such as glycolysis [10]. It is thus well-known that many enzymes are regulated by interactions in large regulatory networks [12].

Building upon a previous pilot study on glycolysis [13], here we investigate the connection between interactomics (PPI networks) and metabolomics (metabolic networks) on a more comprehensive scale, focusing on carbohydrate metabolism. While numerous PPIs have been elucidated experimentally in model organisms such as *E. coli*, the significance of most of them remain unclear, given that their binding affinity is largely unknown but they must clearly play a crucial role in metabolic regulation [14]. Therefore, we have focused on protein abundance data and thus stoichiometry, assuming that the impact of an interaction is likely reflected by its stoichiometry.

We have used published protein abundance data for approximately 2600 *E. coli* proteins across different growth conditions, including multiple carbon sources [15].

Not surprisingly, some protein-enzyme interactions (PEIs) involve uncharacterized proteins as binding partners that co-occur in many species. Such potentially conserved interactions are good candidates for future experimental analysis and thus may reveal novel regulators of metabolism [15–17] as conservation of both components of the protein-enzyme interaction might imply a significant physiological role.

Carbohydrate metabolism has been highly conserved in evolution and may thus even illuminate human metabolic disorders like obesity and diabetes [18–24]. Similarly, an understanding of the metabolome-interactome interface may contribute to applications such as biofuel production [25–27], antibiotic development [11,28] or other applications of metabolism. In fact, a protein-enzyme interaction can be considered to be more widely important if it occurs in multiple species other than *E. coli* [29].

## Materials and Methods

### 2.1 Metabolic enzymes and their pathway information

We obtained the list of metabolic genes of carbohydrate metabolism divided into different carbohydrate pathways from KEGG [2] (**Table 1**, for full list see **Table S4**). These genes were mapped onto proteins of *E. coli* K-12 (proteome ID: UP000000625) using UniProt’s ID mapping feature [30]. This resulted in a list of all *E. coli* metabolic enzymes categorized into 15 carbohydrate metabolic pathways.

### 2.2 Protein-enzyme interaction data

A total of 299 binding partners (“interactors”) and 378 interactions of *E. coli* carbohydrate enzymes were obtained from the IntAct database [31–33], listed in **Table S1**. These 378 PEIs are non-spoke expanded interactions which makes sure the enzyme and protein actually bind to each other. Also, we did not include PPI data from Arifuzzaman et al., 2006 as they used a pull down method to report PPIs which may give false positive result sometimes [34].

### 2.3 Inferring what kind of proteins bind metabolic enzymes

Interactors were classified based on 1) if they are enzymes 2) if they are a metabolic protein and 3) if they are (un-)characterized. UniProt provided the EC number for proteins which are enzymes [35]. We used this feature to classify the interactors into enzymes and non-enzymes first. Then ToppGene (a gene enrichment software [36]) was used to classify the enzyme and non-enzyme interactors into metabolic and non-metabolic (**Fig 1**). The list of interactors was then processed in ToppGene to produce a list of Gene Ontology (GO) Biological Process terms along with the interactors falling under that GO term [37]. Only the “Metabolic Process” GO terms were chosen. If the interactor listed in these GO terms was an enzyme, it was called 1) metabolic enzyme; the remaining enzymes were called 2) non-metabolic enzymes (**Fig 1,2**). In the same way, we classified the non-enzyme interactors into 3) metabolic and 4) non-metabolic proteins. Interactors whose protein names mention “Uncharacterized” or lacked GO annotations were collectively called “unknown proteins”.

**Fig 1.**
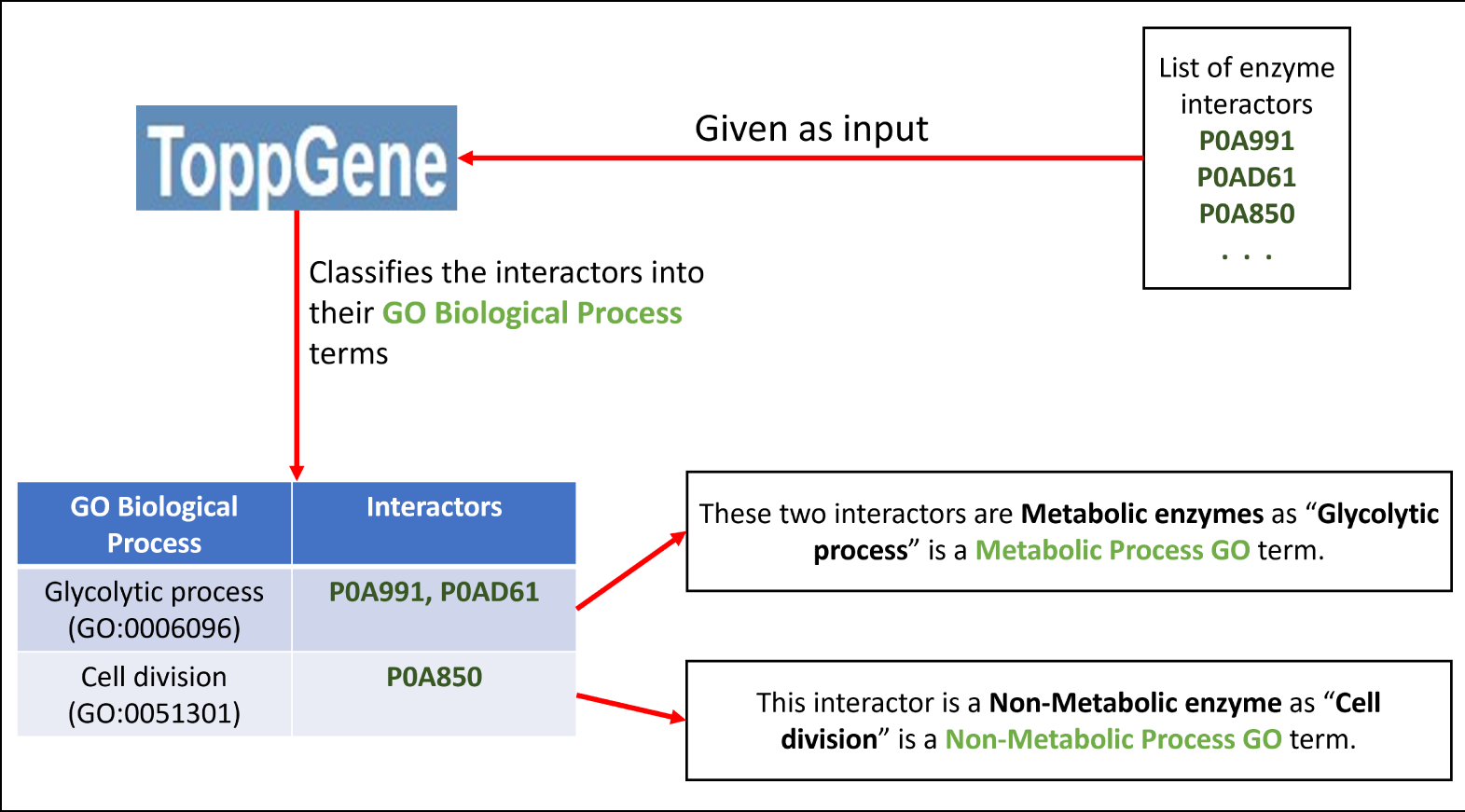
Schematic representation of classifying enzyme interactors into metabolic and non-metabolic.

**Fig 2.**
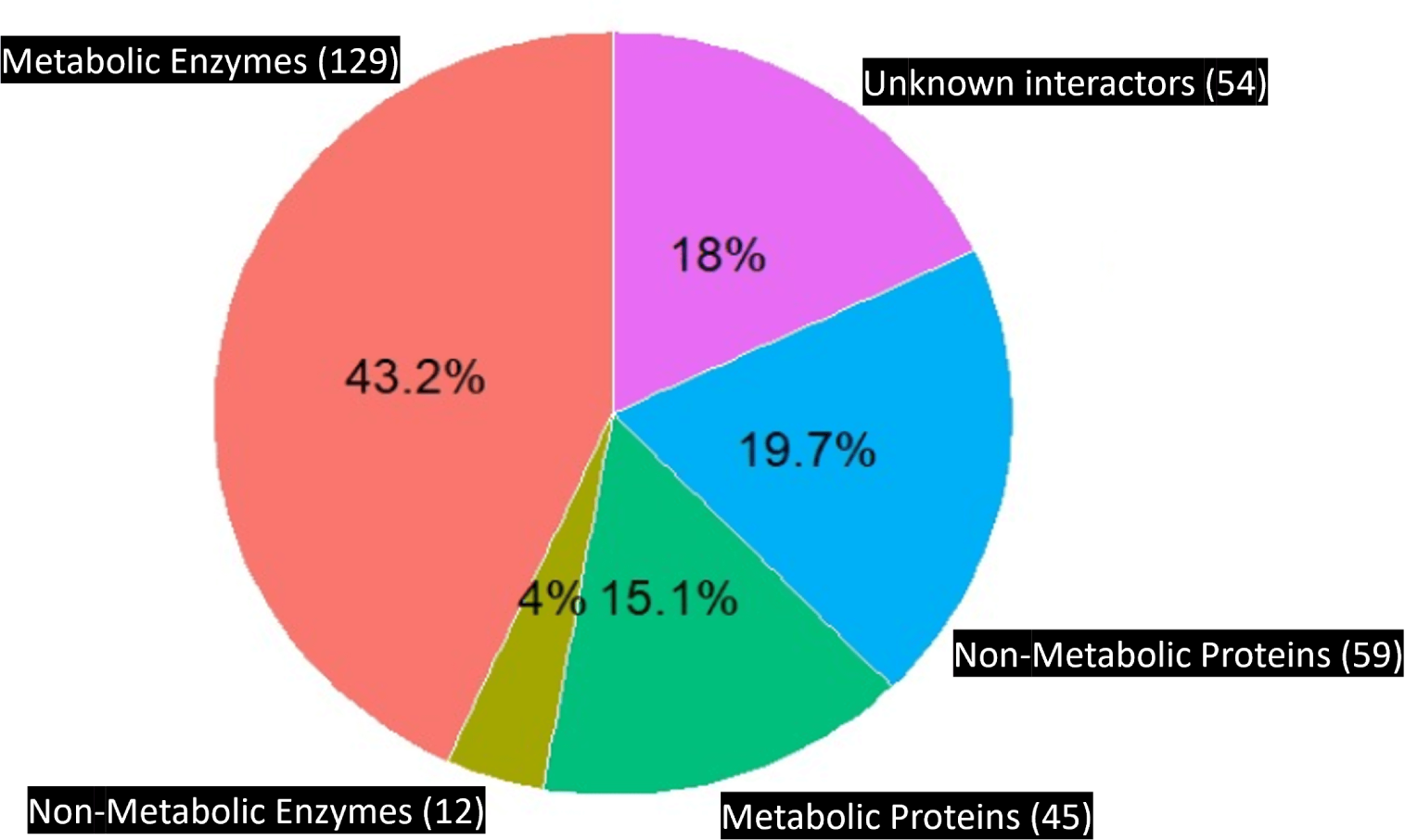
*E. coli* carbohydrate enzymes primarily interact with other metabolic enzymes. They also interact with some non-enzymes comprising 45 metabolic and 59 non-metabolic proteins. Apart from that, *E. coli* carbohydrate enzymes also bind 54 proteins of unknown function.

**Fig 3.**
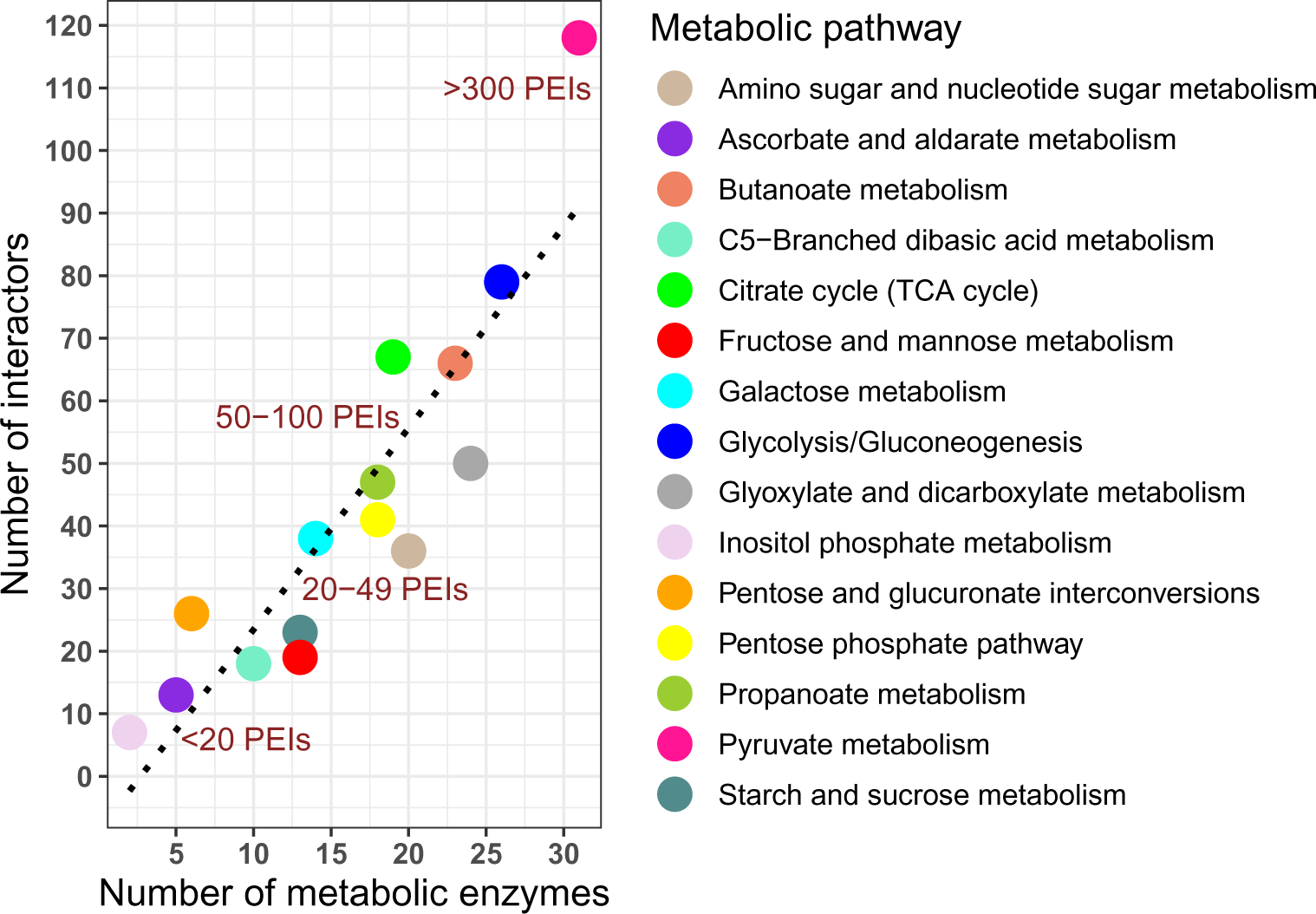
Interactions in *E. coli* carbohydrate pathways correlate with pathway size. Pathways with more enzymes bind more proteins compared to pathways with low count of enzymes.

### 2.4 Conservation of *E. coli* PEIs

The EggNog database provides a list of bacterial ortholog (OG) species for each protein in UniProt [38]. We obtained that list of OG species from EggNOG for all *E. coli* enzymes and their interactors. Presence of the orthologs of both the enzyme and interactor from *E. coli* is likely to indicate a conserved interaction in the ortholog species [29,39]. Based on this, we counted the number of species in which both enzymes and their interactors for all *E. coli* PEIs occur and we called this number “conservation value” (**Fig 4A**). We have also listed some bacterial species known to cause diseases in **Table 6** where many *E. coli* PEIs (or rather predicted PEIs) are conserved.

**Fig 4.**
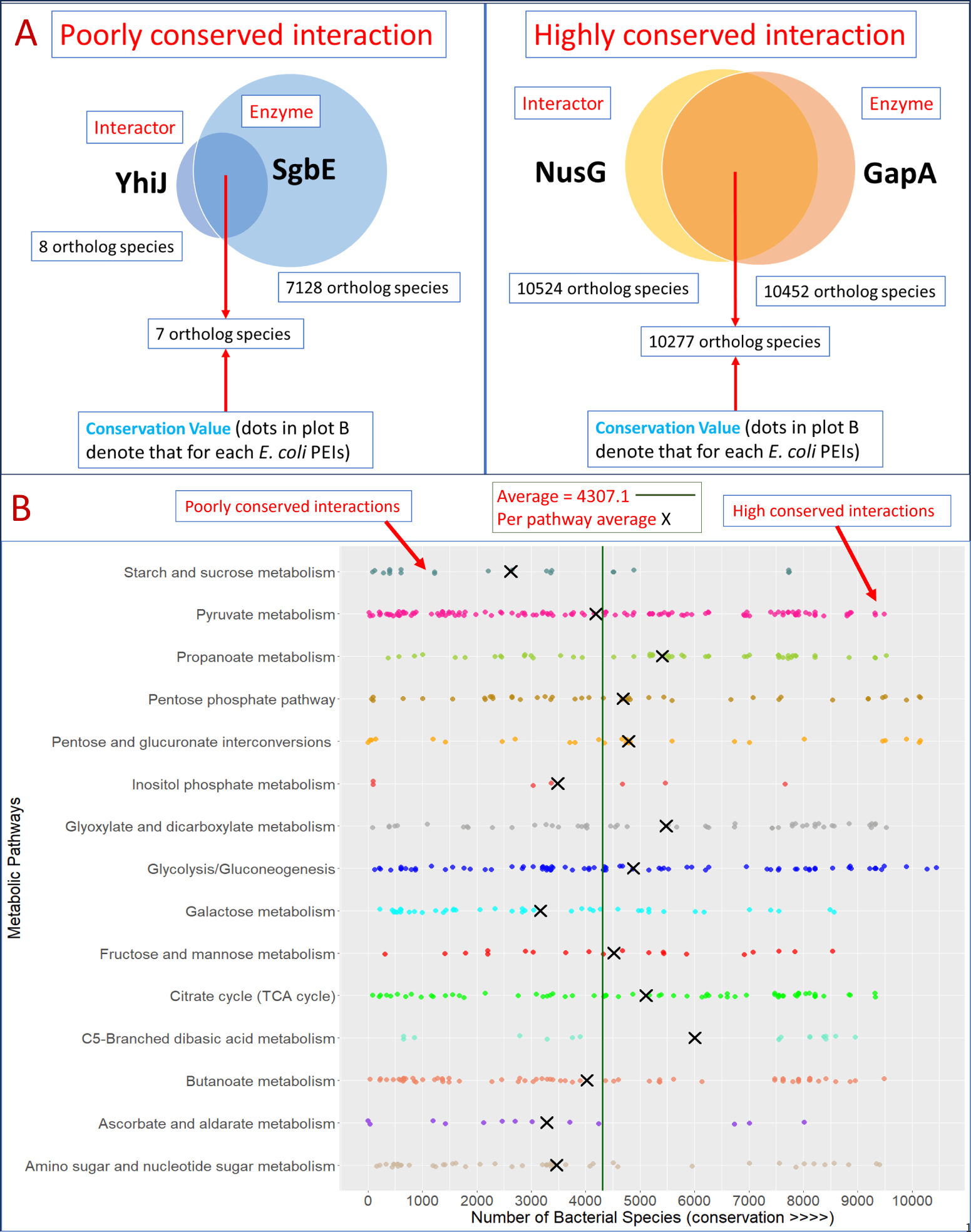
Conservation of interactions in CHM. (A) Examples of highly and poorly conserved PEIs and how they were counted (B) *E. coli* pathways including central carbohydrate pathways have more conserved PEIs than others. The average conservation value is indicated by a vertical line.

### 2.5 Examining protein abundance of *E. coli* protein-enzyme interactions

Schmidt et al. computed abundance of *E. coli* proteins in not 1 or 2 but in 22 different growth conditions [15]. Therefore, we found this study appropriate to test how amounts of PEIs differ when *E. coli* metabolic state changes. Among the total number of *E. coli* proteins found in our dataset, this study provided abundance data in the form of protein copies per cell for approximately 51% of them in 22 different growth conditions (**Fig 5, x axis**). We used this data to obtain and compare abundance values for both enzyme and their binding proteins (**Fig 5, y axis**) across these 22 conditions (**Fig 5,7,8**).

**Fig 5.**
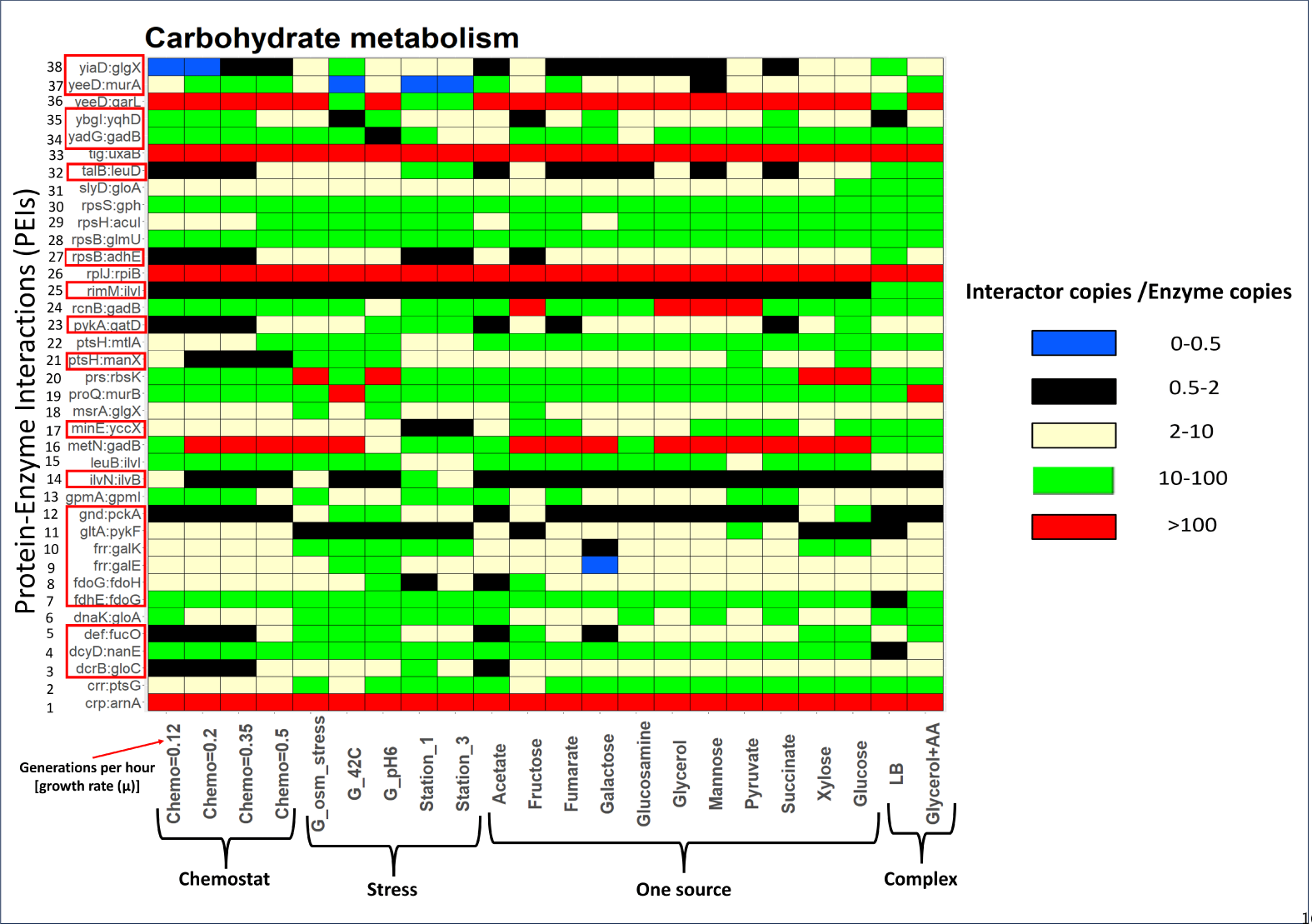
Impact of metabolic state on abundance and ratios of PEIs. The ratio (abundance of interactor/abundance of enzyme) for 38 *E. coli* PEIs (**y axis**) in 22 different *E. coli* environments (**x axis**). Ratios were divided into 5 different ranges shown using a colored brick. For example, blue bricks denote a ratio of 0 - 0.5 while a green brick shows values from 10 - 100. A blue or black brick indicates that the enzyme is more or equally abundant than the interactor in that media. A yellow, green or red brick shows that the interactor is more abundant than the enzyme in that media.

**Fig 6.**
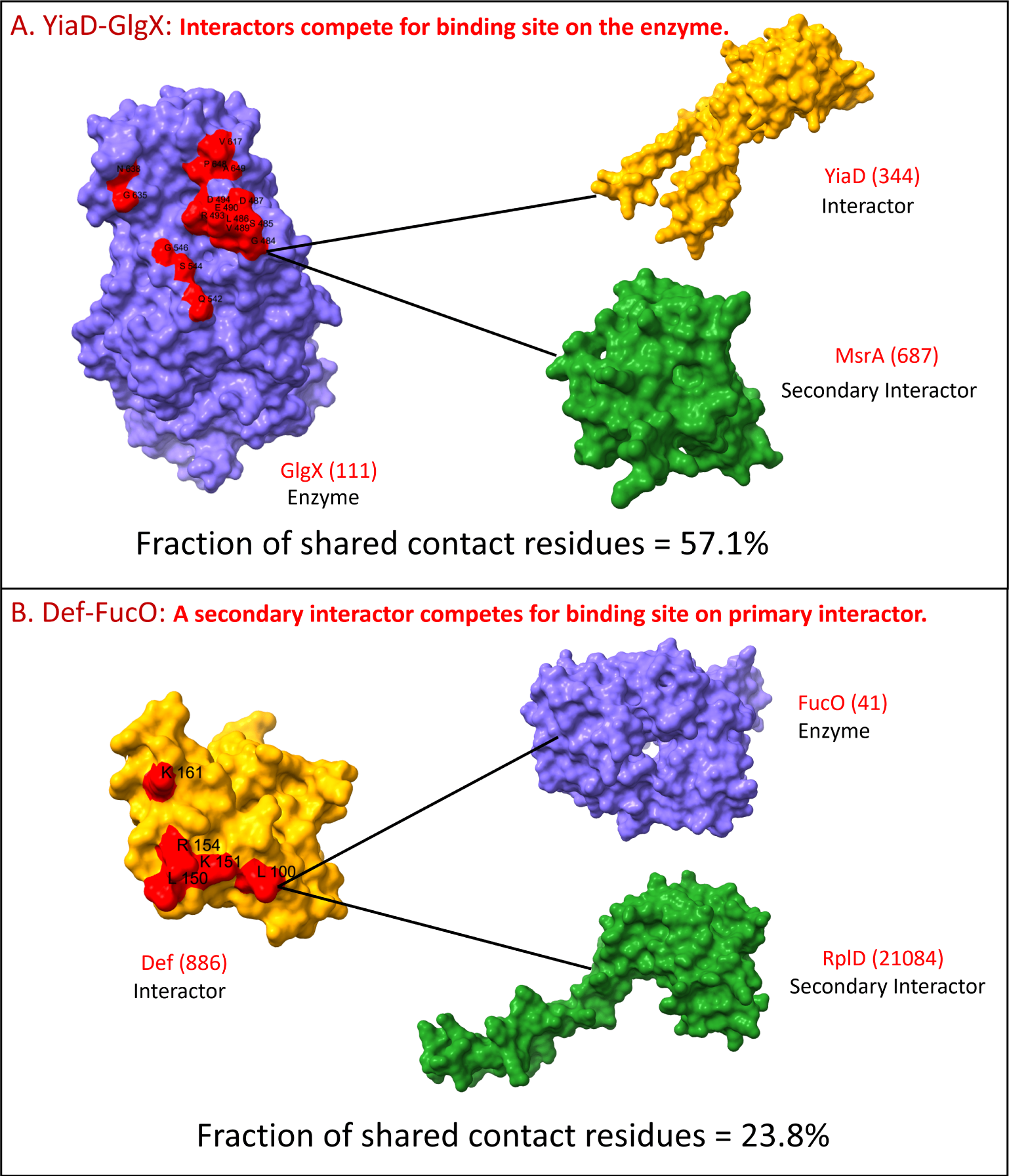
Primary and secondary interactors may compete for the same binding sites. (A) Secondary protein MsrA uses 57% of the binding residues involved in YiaD-GlgX interaction. (B) 24% of Def’s residues binding enzyme FucO are also used by ribosomal protein RplD to bind Def. Def residues binding both FucO and RplD and GlgX residues binding both MsrA and YiaD are shown in red in their protein structures.

**Fig 7.**
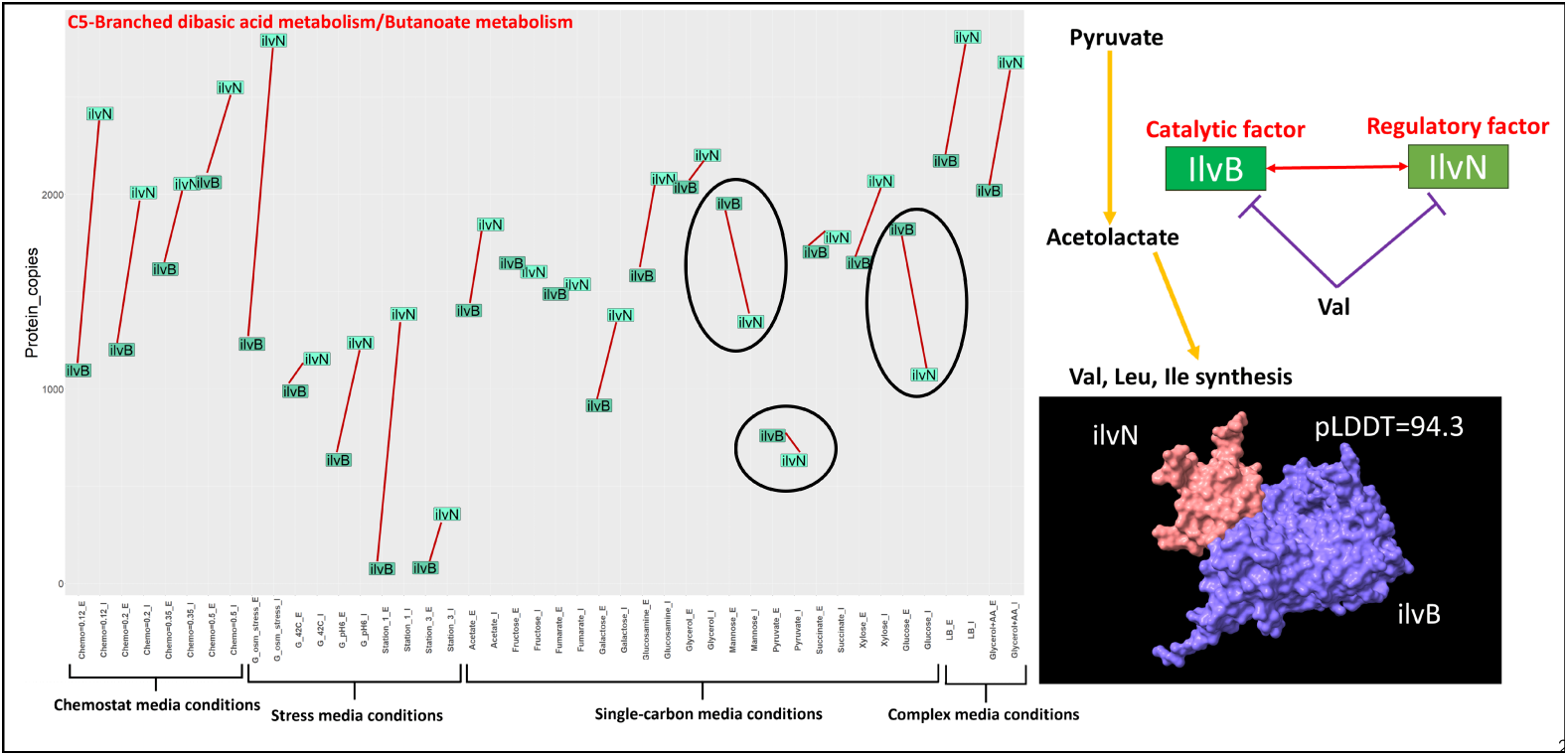
Enzyme IlvB (butanoate and C5-branched dibasic acid metabolism) depends on its interactor IlvN for its function. *E. coli* regulates amino acids production by interaction between IlvN and IlvB as IlvN works as an allosteric factor of IlvB [49–51]. AlphaFold [52] provided structural model of this PEI with a high confidence score (pLDDT = 94.3). Inverse stoichiometry in certain conditions is highlighted in **black** circles.

### 2.6 Visualizing carbohydrate metabolism using KEGG mapper [2]

In order to put carbohydrate metabolism in a broader context, we highlighted CHM in *E. coli*’s metabolome (KEGG: map01100) in **Fig 9**. The figure was constructed using KEGG mapper and each of *E. coli*’s 15 different carbohydrate metabolic pathways were color-coded separately.

**Fig 8.**
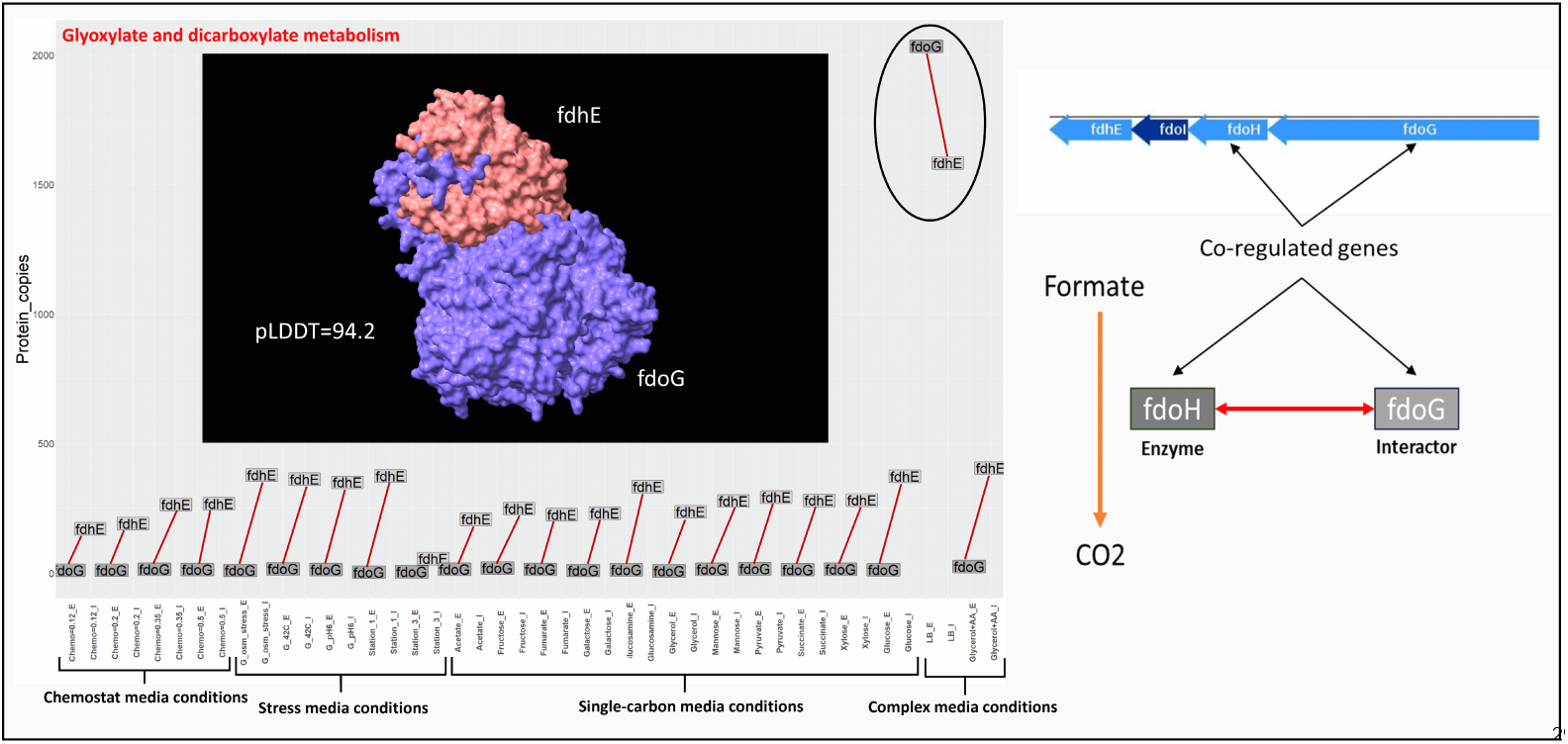
Enzyme FdoG, (dehydrogenizing formate) and its interactor, FdhE simultaneously increase dramatically in LB but are less abundant in other growth conditions. FdhE and FdoG are co-regulated genes [56] and FdhE regulates FdoG when it binds it [55]. AlphaFold [52] predicted the structure of this PEI with high confidence (pLDDT = 94.2). Inversing of stoichiometry in certain conditions are highlighted in **black** circles.

**Fig 9.**
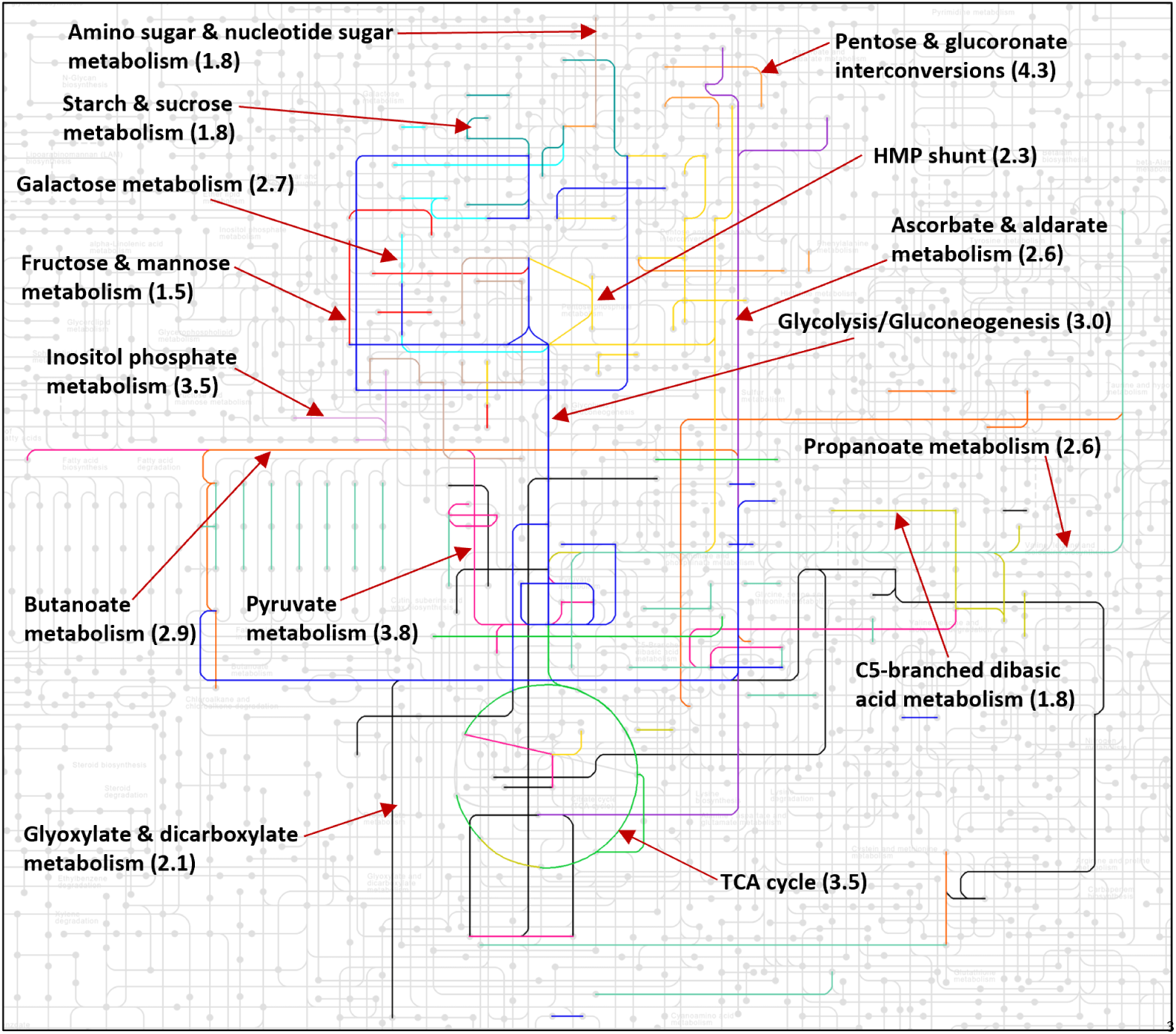
*E. coli* carbohydrate pathway is the biggest of all pathways. With 15 different sub-systems (pathways), we found that *E. coli*’s carbohydrate metabolic pathway has a large number of enzymes (146), proteins (299) and protein-enzyme interactions (378) compared to other pathways such as lipid (33,108,119) or amino acid metabolism (111,200,261) [3,68]. The number in parentheses are average interactor counts per enzyme for that pathway. The *E. coli* metabolome map can be found at https://www.genome.jp/pathway/map01100.

## Results

There are 242 carbohydrate metabolism enzymes in total (**Table 1**). Out of those, there are 146 enzymes (60%) which interact with other proteins (or interactors). There are 299 interactors. A summary of the enzymes and interactors is provided in **Table 2**.

**Table 2.** Summary statistics for metabolic pathways, enzymes, and their interactions in *Escherichia coli*.

| Component | Count |
| --- | --- |
| Carbohydrate metabolic pathways | 15 |
| Carbohydrate metabolic enzymes | 242 |
| Carbohydrate enzymes binding interactors | 146 |
| Interactors binding carbohydrate enzymes | 299 |
| Carbohydrate metabolism protein-enzyme interactions (PEIs) | 378 |

### 3.1 The *E. coli* interactome of carbohydrate metabolism

#### 3.1.1 *Escherichia coli* carbohydrate enzymes bind different types of proteins

While focusing on enzymes associated with carbohydrate metabolism, we first sought to understand the physiological roles of the interactor proteins as they themselves may have varying cellular functions (enzymatic or non-enzymatic and metabolic or non-metabolic). The majority of the interactors were found to be metabolic enzymes (43.2%) (**Fig 2**) [40]. About 35% of the interactors are non-enzymatic proteins and some of them play non-metabolic roles too. Carbohydrate enzymes also bind 12 enzymes having non-metabolic functions (**Fig 2**). Interestingly, a significant number of interactors are unknown proteins with 27 (9%) being uncharacterized and another 27 (9%) lacking annotation [41].

#### 3.1.2 Number of PEIs scale with pathway size

We first classified the *E. coli* protein-enzyme interactions (PEIs) into different carbohydrate metabolic pathways. We found that pathways with more metabolic enzymes bind more proteins and thus, have more PEIs (**Fig 3**). For instance, pyruvate metabolism has *>* 100 PEIs (**Fig 3**).

### 3.2 Conservation of carbohydrate PEIs

We analyzed if *E. coli* PEIs are conserved at the species level [42]. We computed the number of ortholog members between the protein and its binding enzyme for all PEIs to produce a conservation value (**Fig 4A**).

The conservation value of each PEI is represented by a dot in **Fig 4B** and is color coded according to the pathway of the enzyme (**Table 1**). The average conservation value is **4307** meaning that on average 4307 other species (genomes) also have both the enzyme and the interactor of a specific PEI. The average conservation value per pathway (X) is higher than **4307** for most pathways, such as glycolysis or propanoate metabolism. These pathways have more conserved PEIs.

### 3.3 Abundances of metabolic enzymes and their binding partners suggest novel ways of metabolic regulation

#### 3.3.1 The abundance of metabolic enzymes and their interactors change when the metabolic state changes

The stoichiometry of enzymes and their regulators is initially important for metabolic regulation [43]. For instance, a highly abundant inhibitor can essentially block its target enzyme even if their binding affinity is relatively low [43]. Schmidt et al [15] provided abundance data of approx. 2600 *E. coli* proteins in 22 different cell growth conditions (**Fig 5, x axis**). To find the most impactful interactions, we chose the PEIs where the amount of interactor was at least 10 fold higher than its binding enzyme in at least one growth media, that is, the abundance ratio of interactor/enzyme should be *>*= 10 in at least one growth condition. 38 PEIs were found to meet this high interactor to enzyme ratio (**Fig 5**).

First, as expected a protein’s abundance may change when *E. coli*’s growth media changes [44,45] and thereby, the relative abundance (bricks in **Fig 5**) may also change. We found that **PEIs 16, 19, 20, 24 and 36** having interactors at least 10 fold higher than the enzyme in one condition (**green bricks**) became 100 or 1000 fold higher than the enzyme (**red bricks**) in other conditions. There were **5 PEIs (1, 26, 28, 30 and 33)** where interactors/enzyme ratio was constant in all conditions.There were 20 PEIs where concentration of interactors higher in one media than the enzyme become lower than that in another media (ratio *<* 1). These PEIs are **3-5, 7-12, 14, 17, 21, 23, 25, 27, 32, 34, 35, 37 and 38** (**red box** in **Fig 5 y axis**). In these PEI cases, **yellow, green; or red bricks** in one growth media switched to **blue or black brick** in another media. One example is enzyme GlgX, having a lower concentration than YiaD in LB media (PEI 38) (**green brick**) but a higher concentration than YiaD in chemostat cultures (**blue brick**). We assumed these PEIs are good candidates for showing regulatory behavior through binding and hence, were further analyzed.

A change in ratio from one condition to another condition for a PEI can be due to change in abundance of either enzyme or interactor or both. Therefore, we compared absolute abundance of protein and its interactor for these 20 PEIs across 22 different cell media (**Fig S1-S5** and **Fig 7,8**).

A well-known example is PEIs 9 and 10 which involve the enzymes GalK and GalE which increase their abundance in galactose [46] (their concentrations are usually low in other cell conditions) and Frr which maintains a consistent expression level in all cell media conditions implying that it likely does not influence the expression of GalK/GalE (**Fig S1,S2**). We conclude that GalK/E are not regulated by their interactor Frr: if GalK and GalE are expressed at high levels, Frr is not abundant enough and if their levels are low, they are not required for galactose metabolism, even if Frr may be at sufficient level to regulate them.

Similarly, GadB expression is upregulated when *E. coli* faces acidic stress [47]. GadB’s abundance becomes high in glucose media (PEI 34) at pH6 but is otherwise low in other media while its interactor, YadG is sometimes more or less abundant depending on growth conditions (**Fig S3**). However, it is not less abundant than GadB in any of the 22 growth media. Only during acidic stress, the difference in abundance between GadB and YadG decreases due to GadB’s increase in that condition (**Fig S3**). We would like to point out here that YadG is an uncharacterized protein which is more abundant than a *E. coli* enzyme GadB in all 22 growth conditions.

#### 3.3.2 Secondary interactions

The interactions of enzymes with their interactors can be significantly affected by secondary interactions if a secondary interactor competes for binding with the enzyme or the primary interactor (**Fig 6**). Secondary interactions may be significant if the abundance of a secondary interactor is higher than the abundance of an enzyme or its interactors. Excluding the 3 PEIs as mentioned above among the 20 PEIs (**red box** in **Fig 5, y axis**), the remaining 17 PEIs were tested to see if secondary interactions affect them in glucose (minimal) and LB media (rich) respectively. Out of 17 PEIs, there were 5 PEIs (**Table 3**) which may get affected by secondary interactions, i.e. a secondary interactor is present at higher abundance levels than the interactor, so the former may compete with it to bind the enzyme (**Fig 6**/**Table 3**). One good example of the latter is interaction Def:FucO. Secondary interactors RplD, RplV and RplQ bind Def which is the interactor of enzyme FucO. These ribosomal proteins due to their abundance being higher than FucO may occupy most of the binding sites on Def when they bind it, thus, reducing the chances of FucO binding Def (**Fig 6B**/**Table 3**).

**Table 3.** *E. coli* PEIs possibly regulated by secondary interactors. [G/L: value (abundance in glucose/LB media].

| Enzyme (Abundance) | Interactor (Abundance) | Secondary interactors of Enzyme affecting PEI binding | Secondary interactor (Abundance) | Secondary interactors of Interactor affecting PEI binding | Secondary interactor (Abundance) |
| --- | --- | --- | --- | --- | --- |
| GlgX (G: 111) | YiaD (G: 344) | Yes | MsrA (G: 687) | No | EpmB (G: 53/L: 89) |
| IlvI (G: 441/L: 114) | RimM (G: 776/L: 1786) | Yes | IlvH (G: 1533), MqsA (G: 34), LeuB (G: 4588) | Yes | ProQ (G: 2318/L: 4624), IlvF (G: 220/L: 67), RpsS (G: 19008/L: 51051), UbiH (G: 63/L: 114) |
| FdoH (G: 4) | FdoG (G: 13) | No | N/A | Yes | FdhE (G: 374) |
| FucO (G: 41/L: 234) | Def (G: 886/L: 1542) | No | N/A | Yes | Usg (G: 456/L: 583), UbiF (G: 31/L: 46), TrmH (G: 25/L: 70), RplD (G: 21084/L: 49874), RplQ (G: 18398/L: 46700), RplV (G: 49717/L: 137404) |
| GloC (G: 842) | DcrB (G: 2176) | No | N/A | Yes | OsmY (G: 10490) |

**Table 4:** Secondary proteins-affected PEIs.

| Protein-Enzyme Interaction (PEI) | Secondary Interactor binding to Enzyme | Secondary Interactor binding to Interactor | Fraction of overlapping contact residues | Media |
| --- | --- | --- | --- | --- |
| YiaD-GlgX | MsrA | N/A | 57.1% | Glucose |
| RimM-IlvI | LeuB | RpsS | 61.3%/100% | Glucose/LB |
| FdoG-FdoH | N/A | FdhE | 16.3% | Glucose |
| Def-FucO | N/A | RplD | 23.8% | Glucose/LB |
| DcrB-GloC | N/A | OsmY | 42.9% | Glucose |

Among 17 PEIs, these 5 PEIs may get affected by secondary interactors either in glucose or LB or in both. The remaining 12 PEIs, after comparing abundance of secondary interactors with either enzyme or interactor, were predicted to not get affected by secondary interactors (**Section 3.4/Table 5**).

**Table 5.**
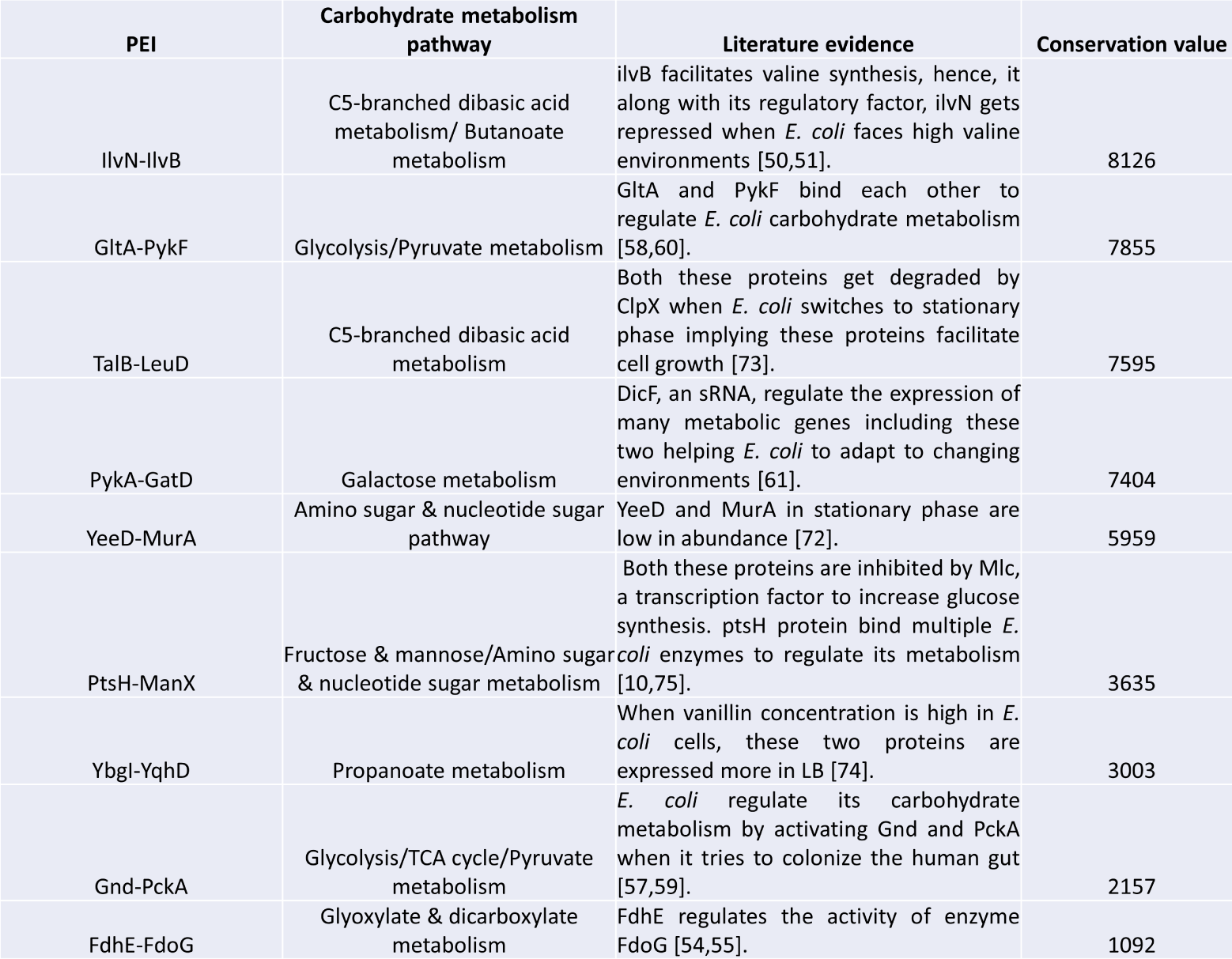
List of potentially regulatory protein-enzyme interactions in *E. coli* metabolism.

**Table 6.**
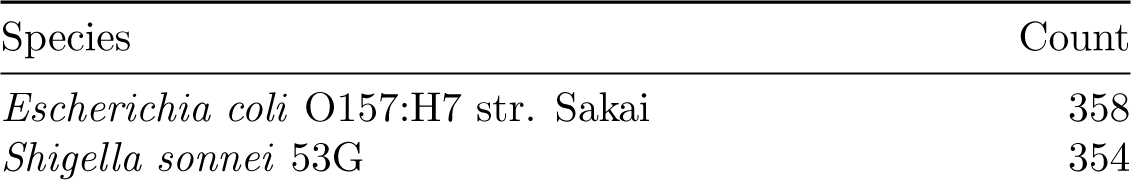
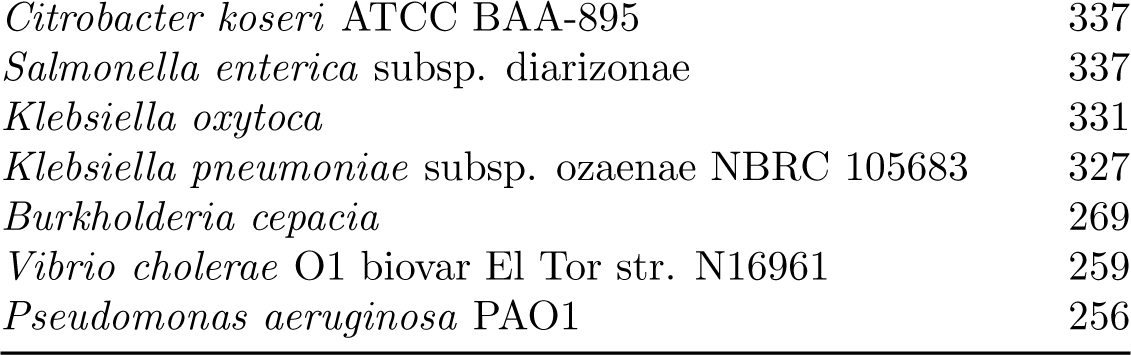
Pathogen species where *E. coli* PEIs co-occur.

| Species | Count |
| --- | --- |
| *Escherichia coli* O157:H7 str. Sakai | 358 |
| *Shigella sonnei* 53G | 354 |
| *Citrobacter koseri* ATCC BAA-895 | 337 |
| *Salmonella enterica* subsp. diarizonae | 337 |
| *Klebsiella oxytoca* | 331 |
| *Klebsiella pneumoniae* subsp. ozaenae NBRC 105683 | 327 |
| *Burkholderia cepacia* | 269 |
| *Vibrio cholerae* O1 biovar El Tor str. N16961 | 259 |
| *Pseudomonas aeruginosa* PAO1 | 256 |

#### 3.3.3 Structural considerations: binding sites

Although secondary interactors may have an impact on the aforementioned interactions, their effect also depends on whether the binding sites of the main interactor and any secondary interactor overlap on an enzyme (**column 4** in **Table 3**). Similarly, the binding sites of a secondary interactor or binding sites of secondary interactor and enzyme overlap on an interactor (**column 7** in **Table 3**). Hence, we calculated what fraction of contact residues of a PEI (main interaction) are likely involved when they bind secondary proteins. We provide those numbers for the “secondary interactor-affecting PEIs” in **Table 4**.

Interactor RimM is a ribosomal protein known to bind many highly abundant proteins [48] including RpsS (**Table 3**. All the RimM residues binding enzyme IlvI also bind secondary interactor RpsS (**Table 4**) whose abundance is ~25 times more than RimM. RimM is more abundant than enzyme IlvI but its secondary interactor RpsS likely occupies all RimM binding sites leaving no space for enzyme IlvI to bind its interactor.

### 3.4 Predicting potential regulators of *Escherichia coli* carbohydrate metabolism

#### 3.4.1 Metabolic regulation by protein-enzyme interactions is confirmed by its published literature

For 9 out of 12 PEIs (**Table 5**), we found published evidence for a regulatory role, hence these 9 *E. coli* PEIs support our approach to predict the ability of PEIs to regulate carbohydrate metabolism (PEIs in red in **Table 5**). The 3 PEIs for which we did not find literature evidence were RpsB-AdhE, DcyD-NanE and MinE-YccX. We also provide conservation values of the 9 PEIs (see **section 3.2**).

### 3.4.2 Case studies of 2 PEIs predicted as metabolism regulators

Enzyme IlvB synthesizes acetolactate from pyruvate which produces the branched chain amino acids valine, leucine and isoleucine [49]. In most growth conditions, IlvB is less abundant than IlvN, however, the abundance of IlvB is comparable to IlvN when *E. coli* is grown in the presence of fructose, pyruvate, succinate or fumarate (**Fig 7**). Mitra et al [50] studied this PEI and found that interactor IlvN allosterically regulates enzyme IlvB by activating its enzyme activity. However, when valine is in excess, it changes the confirmation of IlvN-IlvB interaction such that IlvN inhibits the enzyme activity of IlvB instead of activating it [51].

Enzyme FdoG is the metabolism enzyme [53] and protein FdhE is its binding partner. We found that abundance of FdoG increases when *E. coli* is grown on LB (rich media) (**Fig 8**). Tao et al [54] has shown that FdoG does increase its expression in rich media but not in minimal media. Interestingly, its binding partner FdhE shows similar behavior. It also increases its abundance in LB (**Fig 8**). This shows FdoG may depend on FdhE for its function, hence we predict that FdhE acts as a regulatory factor of FdoG [55]. Also, these two are co-regulated genes found in the same operon [56]. Therefore, FdoG’s activity can get affected by its binding regulator, FdhE especially when *E. coli* needs to metabolize formate.

## Discussion

### *E. coli* carbohydrate enzymes bind other metabolic enzymes the most

The protein partners of *E. coli* CHM enzymes were primarily metabolic enzymes (129 interactors or 43% of all interactors, **Fig 2**). Also, different metabolic enzymes interact with each other to control and coordinate *E. coli*’s carbohydrate metabolism [57–61]. Out of 9 PEIs we predicted as regulators (**Table 5**), three of them, GltA-PykF, PykA-GatD and Gnd-PckA are actually enzyme-enzyme interactions. Carbohydrate enzymes also bind 12 non-metabolic enzymes and 59 non-metabolic proteins which indicates that metabolic enzymes can play moonlighting roles by interacting with non-metabolic partners (**Fig 2**) [62]. Portillo et al has explained how methionine and homocysteine enzymes bind to different proteins to help them in apoptosis, RNA processing, gene regulation, cell growth etc. [63]. Surprisingly, *E. coli* carbohydrate enzymes also interact with 54 unknown proteins of which 3 show potentially relevant expression levels: YadG, YeeD and DcrB. These three unknown proteins are at least 3 times more abundant than their enzyme partners GadB, MurA and GloC respectively. Investigating these interactions involving unknown proteins experimentally may shed light on their metabolic or non-metabolic roles [41,64,65].

### Central carbohydrate pathways are more conserved and have more enzymes, interactors compared to other carbohydrate (non-central) pathways

*Escherichia coli* has 15 carbohydrate pathways which are active in different metabolic states (**Fig 9**). For instance, galactose metabolism is active when the sugar source is galactose [66]. But no matter what the sugar source is, *E. coli* has to use some enzymes of central carbohydrate pathways to metabolize sugar and produce ATP [3]. That is why these pathways are also called primary fueling pathways. These pathways are conserved in many other species (**Fig 4B**). They also have a high number of enzymes, interactors and thus PEIs (**Fig 3**). All pathways span a large range of conservation, i.e. some may be highly conserved in one species but not in others. This is not surprising, given that closely related species will have higher conservation levels than distantly related, ecologically divergent species. Pathways which *E. coli* uses only in specific metabolic environments like starch and sucrose, ascorbate and aldarate, inositol phosphate metabolism and amino sugar and nucleotide sugar metabolism were less conserved (**Fig 4B**) and had fewer enzymes binding a smaller number of proteins (**Fig 3**).

If metabolic regulation happens at protein-protein interaction (PPI) levels, we assume it will happen more for central carbohydrate pathways as they are known to be tightly regulated compared to other metabolism pathways [67] and have large interactomes (**Fig 3**).

Abundance of proteins and their binding enzymes get adjusted according to *E. coli*’s metabolic environment.

We examined the relative abundance of all *E. coli* PEIs and we compared the abundance of enzymes and their binding proteins in 22 different media [15]. In order to have a regulatory effect on the enzyme, interactors have to be present at concentration that are near or exceeding that of enzymes [43], depending on the affinity (which we do not know in most cases). We used this criteria to identify 38 PEIs with interactors being at least 10 fold more abundant than the enzyme in at least one media condition (**Fig 5,10**). Counts of PEIs at 2, 5, 10 fold ratios (interactor copies/enzyme copies) are given in **Fig 10**.

**Fig 10.**
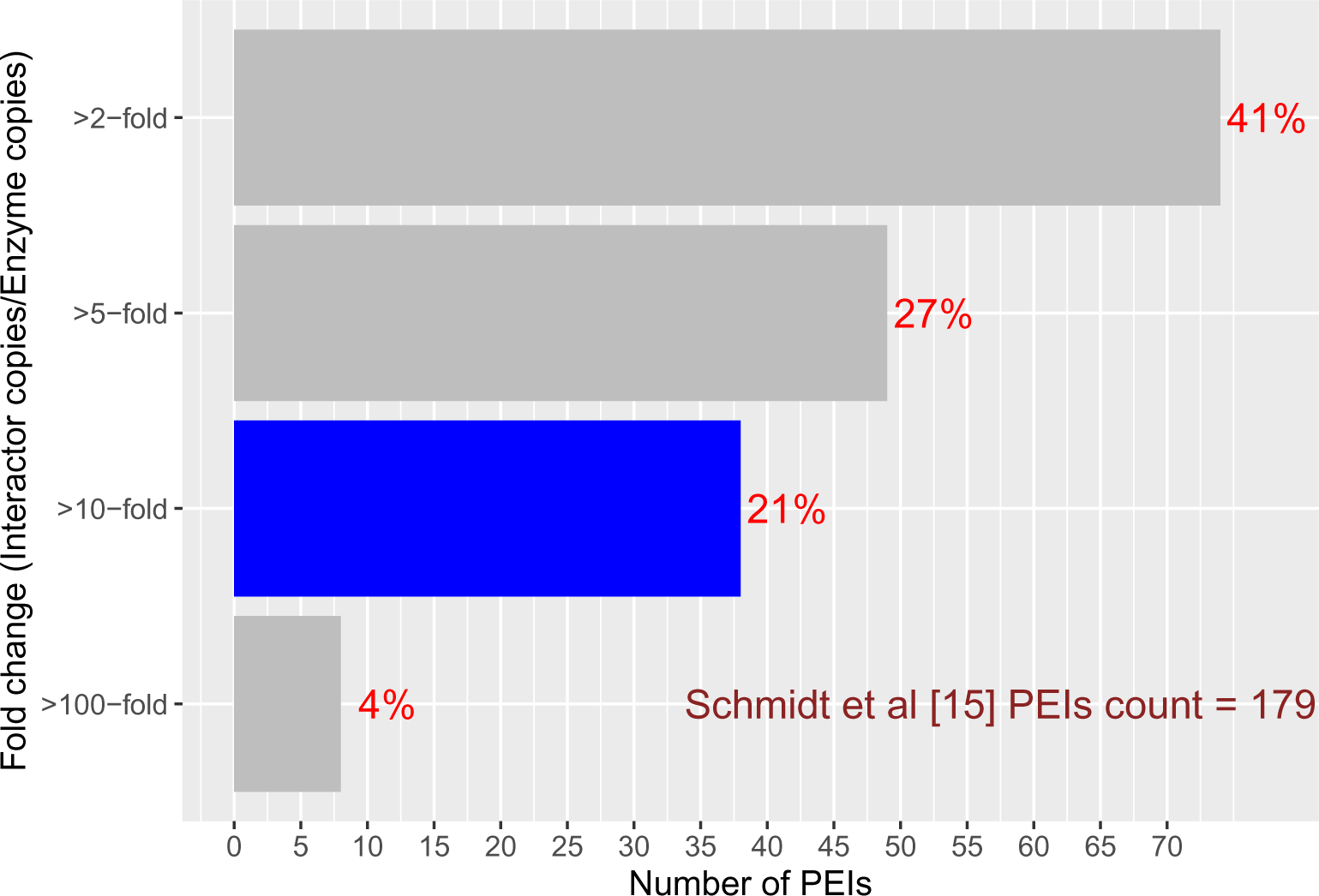
Number of CHM PEIs at specific interactor/enzyme ratios. As fold change increases, number of PEIs decreases. 21% of *E. coli* PEIs (38 PEIs out of 179 PEIs) showed fold change of *>*= 10 in at least one cell growth condition (**blue bar**) [15]. The latter were investigated in detail to delineate their abundance properties in 22 different metabolic states (**Fig 5**) [15].

Interactors that are 10 fold higher than their cognate enzyme in one condition can change their concentration to 0.5 folds, 2 folds, 10, 100 or 1000 folds in other conditions (**Fig 5**). We wondered in which of these cases a 10 fold difference in one condition changes to 0.5 or 2 folds in another condition. This means that *E. coli*’s metabolic condition may affect these PEIs more by increasing enzyme amounts and decreasing interactor amounts which was not the case in another condition. We found 20 *E. coli* PEIs (**red box** in **Fig 5, y axis**) whose relative abundance is not stable but changes substantially when growth conditions change. We also compared absolute abundances of enzymes and interactors in all 22 conditions. For a cell condition, we checked if abundances of enzymes change or interactors change or both change their abundance. We found that the enzymes GalK/GalE and GadB increase their abundances in the interactions GalK/GalE - Frr and GadB - YadG, in galactose and pH stress conditions respectively but their interactors Frr and YadG do not show any condition specific abundances (**Fig S1-S3**). Gal enzymes metabolize galactose [46] and GadB gets expressed more when cells face acidic stress [47]. So, though they interact, we can conclude that these enzymes are not regulated by their interactors in high galactose and high pH media.

### Enzymes and interactors compete with other proteins to bind to each other

Although 20 PEIs (red box in **Fig 5, y axis**) change their relative abundance substantially in certain conditions, we still need to make sure how often do these interactors bind enzymes. For instance, are there other proteins in sufficient amounts to compete with the *E. coli* PEIsWe are calling these proteins secondary interactors (**Fig 6**). We evaluated these based on two hypotheses: (1) the cumulative abundance of secondary interactors is higher than the interactor, so the latter may not get chance to bind the enzyme due to highly abundant competitors. Or (2), cumulative abundance of secondary interactors (of interactor) is much higher than the primary interactor, so that a low abundance enzyme may not get chance to bind the interactor. The 5 PEIs in **Table 3** followed this hypothesis. YiaD-GlgX followed the former hypothesis. FdoG-FdoH, Def-FucO and DcrB-GloC followed the latter hypothesis and RimM-IlvI followed both hypothesis.

However, to prevent primary interactor and enzyme binding, secondary interactors must bind at the sites where these two proteins bind. So, once we had identified their secondary interactor amounts based on above criteria (**Table 3**), we tested if binding site of secondary interactors and enzyme or interactor are likely to overlap (**Table 4**). Binding sites in interactions FucO-Def, FdoH-FdoG, GlgX-YiaD and GloC-DcrB are predicted to overlap with binding sites of secondary interactors, based on threshold distance ChimeraX uses to define two residues in contact [69,70]. **Fig 6** explains two of those scenarios in detail. Protein-protein interactions happen in networks [71] and we tested the network impact by analyzing abundance and binding sites of secondary binders binding either interactor or enzyme. So, though these are one to one protein-enzyme interactions, their other partners can become barriers in these interactions, decreasing the chances of such PEIs to happen in nature. Therefore, this analysis helped us to find relevant *E. coli* PEIs based on secondary interactions (**Table 3,4**).

### *E. coli* protein-enzyme interactions may regulate carbohydrate metabolism

The relevance of our investigation becomes more obvious with specific examples. For instance, the interactor TalB binds the enzyme LeuD and YeeD binds enzyme MurA. The amounts of both the *E. coli* proteins involved in these two interactions decrease in stationary phase [15,72,73] (**Fig S4,S5**), suggesting that these interactions might play a role in promoting growth in *E.coli*. Another example is interactor YqhD which binds YbgI. When *E. coli* cells grown in LB were treated with vanillin, both these proteins were found to be highly abundant [74]. Hence we can assume that vanillin may control this interaction in *E. coli*. A third example involves PtsH which is already known to bind and allosterically regulate multiple *E. coli* metabolic enzymes [10]. Although the latter study do not mention allosteric regulation between PtsH and ManX, there may be one between these two PTS proteins processing glucose. Both are inhibited by a transcription factor, Mlc, when *E. coli* requires more glucose [75]. Thus, PtsH may bind ManX to activate it to facilitate glucose breakdown. Other examples include Gnd - PckA, GltA - PykF and PykA - GatD (**Table 5**) which are enzyme-enzyme interactions known to control and coordinate the carbohydrate metabolism activities [57–61]. IlvN and FdhE are known allosteric factors of IlvB and FdoG enzymes regulating their enzyme activity [50,51,55]. The physiological role of these 9 PEIs could be experimentally tested by measuring enzyme activity with changing stoichiometries which is beyond the scope of this analysis (**Table 5**).

We found thousands of ortholog (OG) pairs of *E. coli* PEIs (**section 3.2**). For example, of the 378 *E. coli* PEIs (**Table 2**) 354 are predicted to co-occur in *Shigella sonnei* 53G. This is not surprising, as *Shigella* and other *γ*-proteobacteria are close relatives of *E. coli*. With increasing phylogenetic distance, the number of orthologous pairs decreases, but equally important, the similarity of proteins also decreases, so many of the predicted interactions based on sequence similarity may be lost in evolution. Nevertheless, we expect that PEIs can regulate metabolism not only in *E. coli* but in many other species where they co-occur. In **Table 10**, we list some pathogenic species where *E. coli* PEIs are predicted to co-occur. Hence, experimental investigation of *E. coli* PEIs may help us to develop novel therapeutic strategies [76] to cure diseases caused by these pathogens.

## Conclusion

This project is the first attempt to look at regulation of carbohydrate metabolism via the binding partners of metabolic enzymes. We predicted some *E. coli* protein-enzyme interactions as potential regulators of its carbohydrate metabolism [28,77]. Out of 299 interactors, the 54 uncharacterized proteins binding *E. coli* carbohydrate enzymes may be especially good candidates for experimentally testing as their biological role is poorly understood [41,65]. Notably, protein-protein interactions are likely to regulate important aspects of human microbiome metabolism [78]. This will drive research into more complex microbiome metabolic interactions including host-pathogen interactions.

